# Rapid functional divergence of grass duplicate genes

**DOI:** 10.1101/490524

**Authors:** Xueyuan Jiang, Raquel Assis

## Abstract

Gene duplication has played an important role in the evolution and domestication of flowering plants. Yet little is known about how plant duplicate genes evolve and are retained over long timescales, particularly those arising from small-scale duplication (SSD) rather than whole-genome duplication (WGD) events. Here we address this question in the Poaceae (grass) family by analyzing gene expression data from nine tissues of *Brachypodium distachyon*, *Oryza sativa japonica* (rice), and *Sorghum bicolor* (sorghum). Consistent with theoretical predictions, expression profiles of most grass genes are conserved after SSD, suggesting that functional conservation is the primary outcome of SSD in grasses. However, we also uncover support for widespread functional divergence, much of which occurs asymmetrically via the process of neofunctionalization. Moreover, neofunctionalization preferentially targets younger (child) duplicate gene copies, is associated with RNA-mediated duplication, and occurs quickly after duplication. Further analysis reveals that functional divergence of SSD-derived genes is positively correlated with both sequence divergence and tissue specificity in all three grass species, and particularly with anther expression in *B. distachyon*. Therefore, as found in many animal species, SSD-derived grass genes often undergo rapid functional divergence that may be driven by natural selection on male-specific phenotypes.

## INTRODUCTION

Angiosperms, or flowering plants, compose one of the most evolutionarily and phenotypically diverse group of eukaryotes. Findings stemming from comparative genomic and experimental studies have led researchers to hypothesize that this extraordinary diversity is primarily a product of gene duplication events (Zhang 2003; Flagel and Wendel 2009; Van de Peer, et al. 2009). For one, duplicate genes are more abundant in angiosperms than in any other sequenced taxonomic group (Zhang 2003; Flagel and Wendel 2009), and differences in numbers of duplicates often contribute to genome sizes that differ by many orders of magnitude, even between closely related species (Flavell, et al. 1974; Bennetzen, et al. 2005). Second, a number of studies have shown that gene duplication can promote the origin of novel plant phenotypes (Flagel and Wendel 2009; Van de Peer, et al. 2009), and that it was likely a key driving factor in the domestication of flowering plants (Hilu 1993; Dubcovsky and Dvorak 2007; Meyer, et al. 2012; Salman-Minkov, et al. 2016). However, many of these findings are associated with studies of duplicates derived from whole-genome duplication (WGD) events, which occurred several times during the past 200 million years of angiosperm evolution (Paterson, et al. 2004; Lockton and Gaut 2005; Cui, et al. 2006; Van de Peer, et al. 2009; Jiao, et al. 2011; Rensing 2014; Panchy, et al. 2016). Yet substantial evidence shows that, in both plants and animals, duplicates deriving from WGD and small-scale duplication (SSD) events differ in quantifiable ways, such as evolutionary rate, essentiality, and function (Hakes, et al. 2007; Carretero-Paulet and Fares 2012; Rensing 2014; Maere and Van de Peer, 2010). Therefore, an open question is how SSD-derived genes in angiosperms evolve and are retained over long evolutionary timescales.

In the simplest case, SSD creates two copies of an ancestral single-copy gene. Considering directionality of duplication, the copy representing the ancestral gene is often called the “parent”, whereas the copy generated by duplication is termed the “child” (Han and Hahn 2009; Assis and Bachtrog 2013). Four mechanisms may underlie the evolution and long-term retention of gene copies in such a scenario. First, under conservation, the ancestral function is preserved in each copy after duplication, perhaps due to a beneficial effect of increased gene dosage (Ohno 1970). Second, under neofunctionalization, one copy preserves the ancestral function, whereas the other copy acquires a new function (Ohno 1970). Third, under subfunctionalization, the ancestral function is divided between copies (Force, et al. 1999; Stoltzfus 1999). Last, under specialization, rapid subfunctionalization is followed by neofunctionalization, resulting in both copies having distinct functions from one another and from their ancestral gene (He and Zhang 2005; Rastogi and Liberles 2005).

Though examples of all of these hypothesized retention mechanisms exist in angiosperms (Duarte, et al. 2005; Throude, et al. 2009; Marcussen, et al. 2010; Bekaert, et al. 2011; Aklilu, et al. 2013; Ma, et al. 2015; Zhang, et al. 2015), their relative abundances on a genome-wide scale remain unknown. One of the reasons for this gap in knowledge is the lack of methods for assessing functional divergence after gene duplication. To overcome this obstacle and distinguish among retention mechanisms of duplicate genes, researchers developed a phylogenetic approach that compares expression profiles between the ancestral single-copy gene in one species and the parent and child copies arising from a SSD event in a closely related sister species (Assis and Bachtrog 2013). Application of their approach to RNA-seq data from two *Drosophila* species suggested that approximately 65% of duplicate genes underwent neofunctionalization (Assis and Bachtrog 2013). Further analyses revealed that neofunctionalization often occurs within a few million years of duplication, results in acquisition of new functions by child copies that arose via RNA-mediated mechanisms, and generates testis-specific gene functions (Assis and Bachtrog 2013; Assis 2014). In contrast, examination of RNA-seq data from eight mammals showed that only 33% of duplicate genes were retained by neofunctionalization (Assis and Bachtrog 2015). The majority of duplicates were instead retained by conservation, and expression divergence was found to occur more gradually in mammals than in *Drosophila*, result in acquisition of new functions equally by parents and children, and generate a diversity of tissue-specific gene functions (Assis and Bachtrog 2015).

Natural selection may act more efficiently in *Drosophila* duplicate genes due to their much larger effective population sizes (*N*_e_) than mammals (Lynch and Conery 2003), which may have contributed to the higher rates of expression divergence observed in *Drosophila* duplicate genes (Assis and Bachtrog 2013, 2015). In particular, the efficiency of selection is proportional to *N*_e_×*s*, where *s* is the selective advantage of a beneficial mutation (Kimura 1983; Charlesworth 2009). Therefore, because angiosperms have comparable *N*_e_ to mammals (Lynch and Conery 2003; Ai et al. 2012; Adugna and Bekele 2015; Stritt et al. 2017, we might expect similar levels of expression conservation between duplicate genes of flowering plants and those of mammals.

In this study, we assess the genome-wide roles of duplicate gene retention mechanisms after SSD in three closely related self-pollinating (Beachell, et al. 1938; Dje, et al. 2000; Gordon, et al. 2014) angiosperms in the Poaceae (grass) family: *Brachypodium distachyon, Oryza sativa japonica* (rice), and *Sorghum bicolor* (sorghum). *B. distachyon* and *O. sativa japonica* share a more recent common ancestor 40-54 million years ago (MYA), and the most recent common ancestor of all three species occurred 45-60 MYA (Bowers, et al. 2005; Bennetzen 2007; Paterson, et al. 2009; International Brachypodium Initiative 2010). Grasses represent an interesting evolutionary system because they are agriculturally important (Davidson, et al. 2012) and, thus, have undergone domestication events in their recent evolutionary histories. Further, these three grass species are ideal for comparison due to the availability of RNA-seq data from the same nine tissues (leaf, anther, endosperm, early inflorescence, emerging inflorescence, pistil, embryo, seed 5 days after pollination, and seed 10 days after pollination) that were obtained in a single lab under similar experimental conditions (Davidson, et al. 2012). Hence, we have a powerful toolkit with which to assess expression divergence after SSD in grasses.

## RESULTS

### Retention mechanisms of SSD-derived duplicates in grasses

A primary goal of our study was to understand how a pair of SSD-derived grass duplicate genes evolves and is retained after its emergence from a single-copy ancestral gene. Therefore, considering the phylogenetic tree depicted in Figure 1, we were interested in pairs of duplicates that arose via SSD along the lineages of *B. distachyon* and *O. sativa japonica* after their divergence from *S. bicolor* (orange stars), *B. distachyon* after its divergence from *O. sativa japonica* (blue stars), *O. sativa japonica* after its divergence from *B. distachyon* (green stars), and *S. bicolor* after its divergence from *B. distachyon* and *O. sativa japonica* (purple stars). To identify such duplicates, we obtained a table of gene family sizes for 16 monocots and their full species phylogeny from the PLAZA 3.0 database (Proost, et al. 2014; Figure S1). Then, we used a maximum likelihood-based approach (Csűös 2010) to ascertain all duplications and losses that occurred along the monocot phylogeny. We applied parsimony rules to identify pairs of duplicates that arose along the branches indicated in Figure 1 (see Figure S1 for full tree). It is important to note that the most recent WGD event in monocots occurred approximately 65 MYA (Jiao, et al. 2011), which is before the divergence of *B. distachyon, O. sativa japonica*, and *Sorghum bicolor*. Therefore, given the size of the monocot tree and number of outgroups considered, the duplications that we extracted with this approach are more likely to be created by SSD rather than WGD events. Next, we required that both duplicate genes, as well as their single-copy ancestral gene in the closer of the two sister species considered, be expressed in at least one tissue (see Materials and Methods for details). This analysis yielded 272 SSD-derived gene pairs in *B. distachyon*, 289 pairs in *O. sativa japonica*, and 340 pairs in *S. bicolor* (Figure 1). Using sequence and synteny information, we inferred the most likely parent and child copy for each pair of duplicates in this dataset (see Materials and Methods for details).

**Figure 1.**
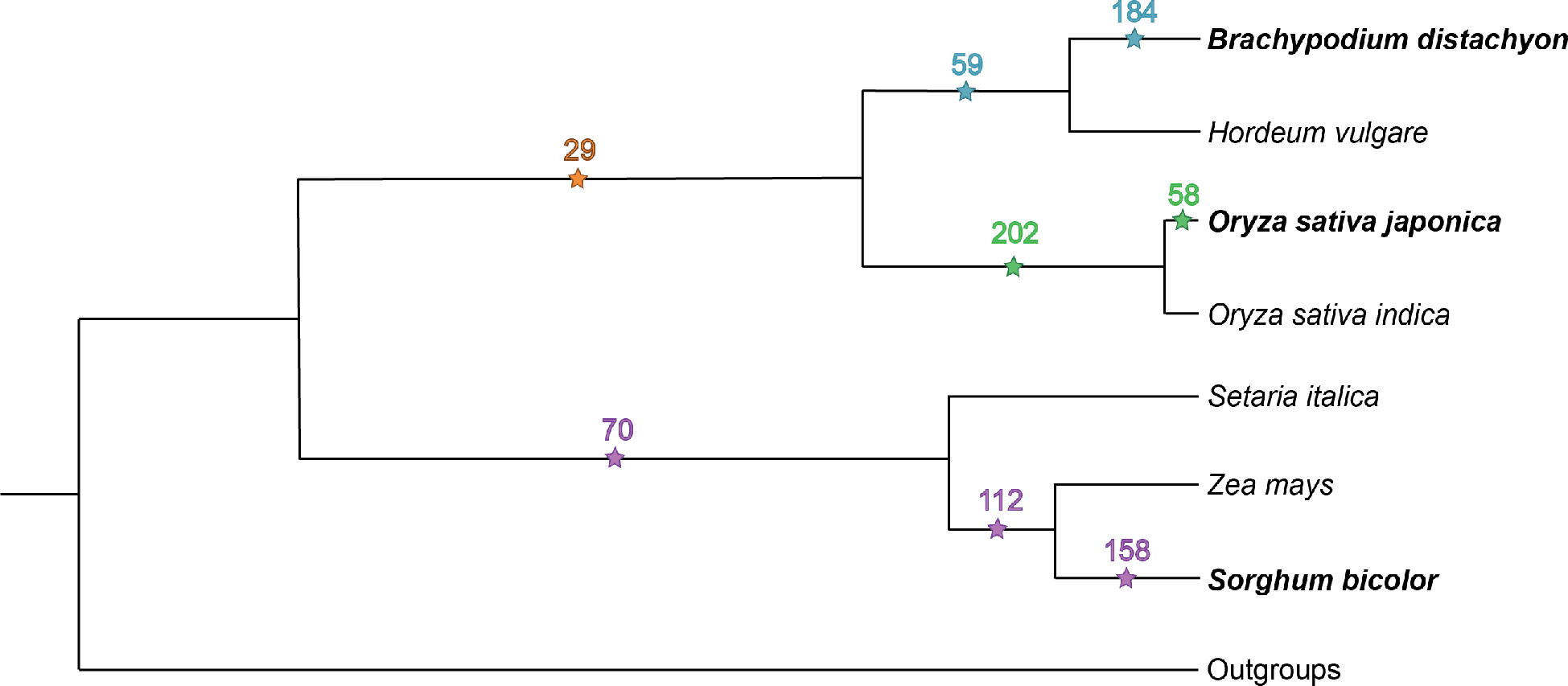
SSD-derived grass duplicate genes ascertained for our analysis. Numbers of SSD-derived duplicate gene pairs that arose along the *B. distachyon* (blue and orange stars), *O. sativa japonica* (green and orange stars), and *S. bicolor* (purple stars) lineages at specified divergence times on the monocot phylogeny. Outgroups used to polarize duplication events were *Musa acuminata, Arabidopsis thaliana, Carica papaya, Populus trichocarpa, Vitis vinifera, Solanum lycopersicum, Physcomitrella patens, Ostreococcus lucimarinus*, and *Chlamydomonas reinhardtii* (see Figure S1 for full phylogeny).

To classify retention mechanisms of SSD-derived grass duplicate genes, we applied the phylogenetic method developed by Assis and Bachtrog (Assis and Bachtrog 2013) to expression profiles constructed from RNA-seq data in nine tissues (Davidson, et al. 2012) of single-copy, ancestral, parent, and child genes of *B. distachyon*, *O. sativa japonica*, and *S. bicolor*. In particular, this method (Assis and Bachtrog 2013) first utilizes the distribution of Euclidian distances between expression profiles of single-copy genes to establish a cutoff that represents the expected expression divergence between two species. Next, it computes Euclidian distances between ancestral and parent expression profiles, ancestral and child expression profiles, and ancestral and combined parent-child expression profiles. Last, it classifies retention mechanisms of each pair of duplicates based on phylogenetic rules. Briefly, the expression profile of the ancestral gene is expected to be similar to those of both the parent and child under conservation, to those of one copy but not the other under neofunctionalization, and to those of neither copy under subfunctionalization or specialization. Distinguishing between subfunctionalization and specialization requires an additional comparison of ancestral and combined parent-child expression profiles. Similarity between these expression profiles suggests that the function of the ancestral gene was subdivided between parent and child copies due to subfunctionalization, whereas dissimilarity points to functional divergence among all three genes due to specialization (Assis and Bachtrog 2013).

Application of the described classification approach (Assis and Bachtrog 2013) uncovered similar proportions of each retention mechanism among *B. distachyon*, *O. sativa japonica*, and *S. bicolor* SSD-derived duplicates (Table 1). Therefore, it appears that genes in all three grass species traverse similar evolutionary paths after SSD. In total, 60.6% of SSD-derived grass duplicates are conserved, 23.8% are neofunctionalized, 0.4% are subfunctionalized, and 15.2% are specialized. Hence, conservation is the most prevalent retention mechanism, indicating that SSD typically results in increased gene dosage in grasses. This level of functional conservation is higher than observed in *Drosophila* (Assis and Bachtrog 2013) and similar to that observed in mammals (Assis and Bachtrog 2015). Thus, our observation is consistent with the smaller *N*_e_ of grass and mammalian species compared with *Drosophila* (Lynch and Conery 2003; Ai et al. 2012; Adugna and Bekele 2015; Stritt et al. 2017).

**Table 1.**
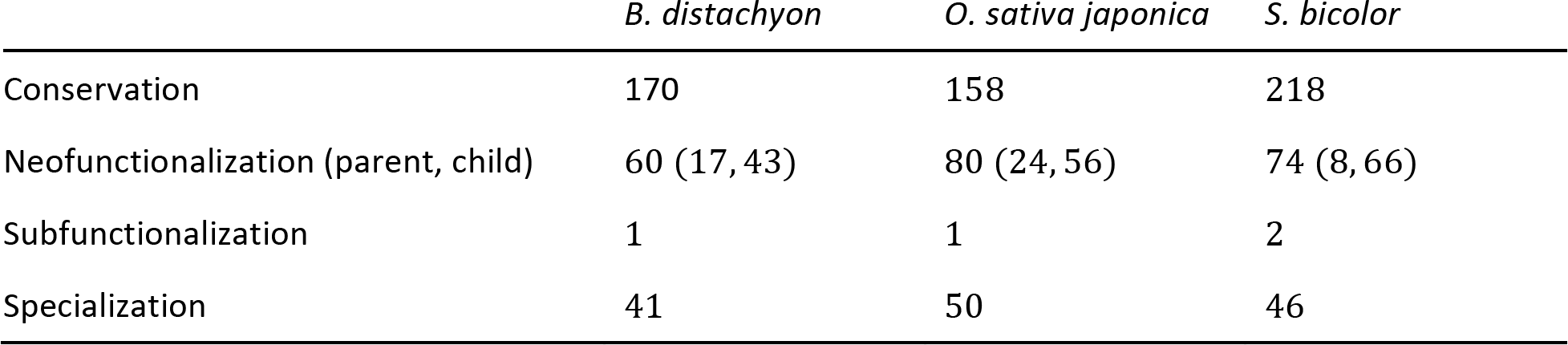
Classified retention mechanisms of SSD-derived grass duplicate genes.

### Contribution of duplication mechanism to expression divergence of SSD-derived grass duplicates

Despite a prominent role of conservation, over one-third of SSD-derived grass duplicate genes undergo expression divergence, most of which occurs asymmetrically via neofunctionalization. This pattern of asymmetric expression divergence is consistent with findings in both *Drosophila* (Assis and Bachtrog 2013) and mammals (Assis and Bachtrog 2015). However, as in *Drosophila* (Assis and Bachtrog 2013) but not mammals (Assis and Bachtrog 2015), neofunctionalization in grasses is also biased in that approximately 72% of *B. distachyon*, 70% of *O. sativa japonica*, and 89% of *S. bicolor* neofunctionalized genes are child copies (Table 1). In *Drosophila*, this bias was associated with RNA-mediated duplication (Assis and Bachtrog 2013; Assis 2014), which produces child copies lacking the introns and regulatory elements of their ancestral genes. The new genomic context of RNA-mediated child duplicates may increase their likelihood of possessing or acquiring novel gene functions (Kaessmann, et al. 2009). Therefore, we hypothesized that RNA-mediated duplication may contribute to biased neofunctionalization of children in grasses as well. To test this hypothesis, we compared observed and expected counts of DNA-and RNA-mediated duplicates retained by conservation, neofunctionalization of parents, neofunctionalization of children, and specialization (Table 2; see Materials and Methods for details). Indeed, there is an overrepresentation of RNA-mediated duplicates retained by neofunctionalization of children (*P* = 0.01, χ^2^ test; see Materials and Methods for details), but not by any other mechanism. This finding indicates that RNA-mediated duplication is more likely to generate children with novel functions in grasses. Moreover, because this pattern exists in both grasses and *Drosophila* (Assis and Bachtrog 2013), it is possible that RNA-mediated duplication acts as a reservoir of functional innovation across many diverse taxonomic groups.

**Table 2.** Observed (expected) DNA-and RNA-mediated SSD-derived duplicates mechanism.

|  | DNA-mediated | RNA-mediated | *P* |
| --- | --- | --- | --- |
| Conservation | 464 (455.98) | 28 (36.02) | 0.17 |
| Neofunctionalization of parent | 39 (38.00) | 2 (3.00) | 0.55 |
| Neofunctionalization of child | 126 (134.39) | 19 (10.61) | 0.01 |
| Specialization | 80 (80.63) | 7 (6.37) | 0.79 |

If RNA-mediated duplication contributes to neofunctionalization in grasses, then we might expect expression divergence to occur either as a byproduct of SSD or soon afterward. Therefore, next we were interested in ascertaining the timing of expression divergence after SSD in grasses. If expression divergence is rapid, then we expect the frequencies of retention mechanisms to be similar among duplicates that arose at different time points in monocot evolution, as was observed in *Drosophila* (Assis and Bachtrog 2013). Alternatively, if expression divergence occurs more gradually after SSD, then we expect higher frequencies of conservation in duplicates that arose more recently and higher frequencies of divergence in those that arose more distantly in the past, as was observed in mammals (Assis and Bachtrog 2015). To address this question in grasses, we divided the duplicates in our dataset into three age classes based on when SSD occurred along the monocot phylogeny and compared observed and expected counts of retention mechanisms in each age class (Tables S1-S3; see Materials and Methods for details). Consistent with findings in *Drosophila* (Assis and Bachtrog 2013), but not in mammals (Assis and Bachtrog 2015), proportions of retention mechanisms are similar among duplicates that arose by SSD at different time points in all three species. Therefore, it appears that functional divergence of SSD-derived grass duplicates often occurs either as a consequence of duplication or shortly afterward.

### Sequence-and tissue-specific correlates with expression divergence of SSD-derived grass duplicates

Previous studies have demonstrated that expression divergence is often positively correlated with protein-coding sequence divergence of duplicate genes in many species (Gu, et al. 2002; Makova and Li 2003; Conant and Wagner 2004; Zhang, et al. 2004; Li, et al. 2005; Assis and Bachtrog 2013; Chau and Goodisman 2017). To assess this relationship in grasses, we calculated Pearson’ s correlation coefficients (*r*) between expression divergence (Euclidian distance) and both nonsynonymous sequence divergence (*K*_a_) and nonsynonymous-to-synonymous sequence divergence (*K*_a_/*K*_s_) rates of each SSD-derived duplicate gene and its ancestral gene in *B. distachyon, O. sativa japonica*, and *S. bicolor* species (Figure 2; see Materials and Methods for details). In all three species, there is a moderately strong positive correlation between expression divergence and *K*_a_ (Figure 2A; *r* = 0.40 − 0.48; *P* < 0.001 for all comparisons, *t* tests; see Materials and Methods for details), and a weak positive correlation between expression divergence and *K*_a_/*K*_s_ (Figure 2B; *r* = 0.10 − 0.17, *P* < 0.05 for all comparisons, *t* tests; see Materials and Methods for details). Thus, expression divergence of SSD-derived duplicates is significantly associated with protein-coding sequence divergence rates, suggesting that expression patterns and encoded proteins of grass duplicate genes evolve in tandem.

**Figure 2.**
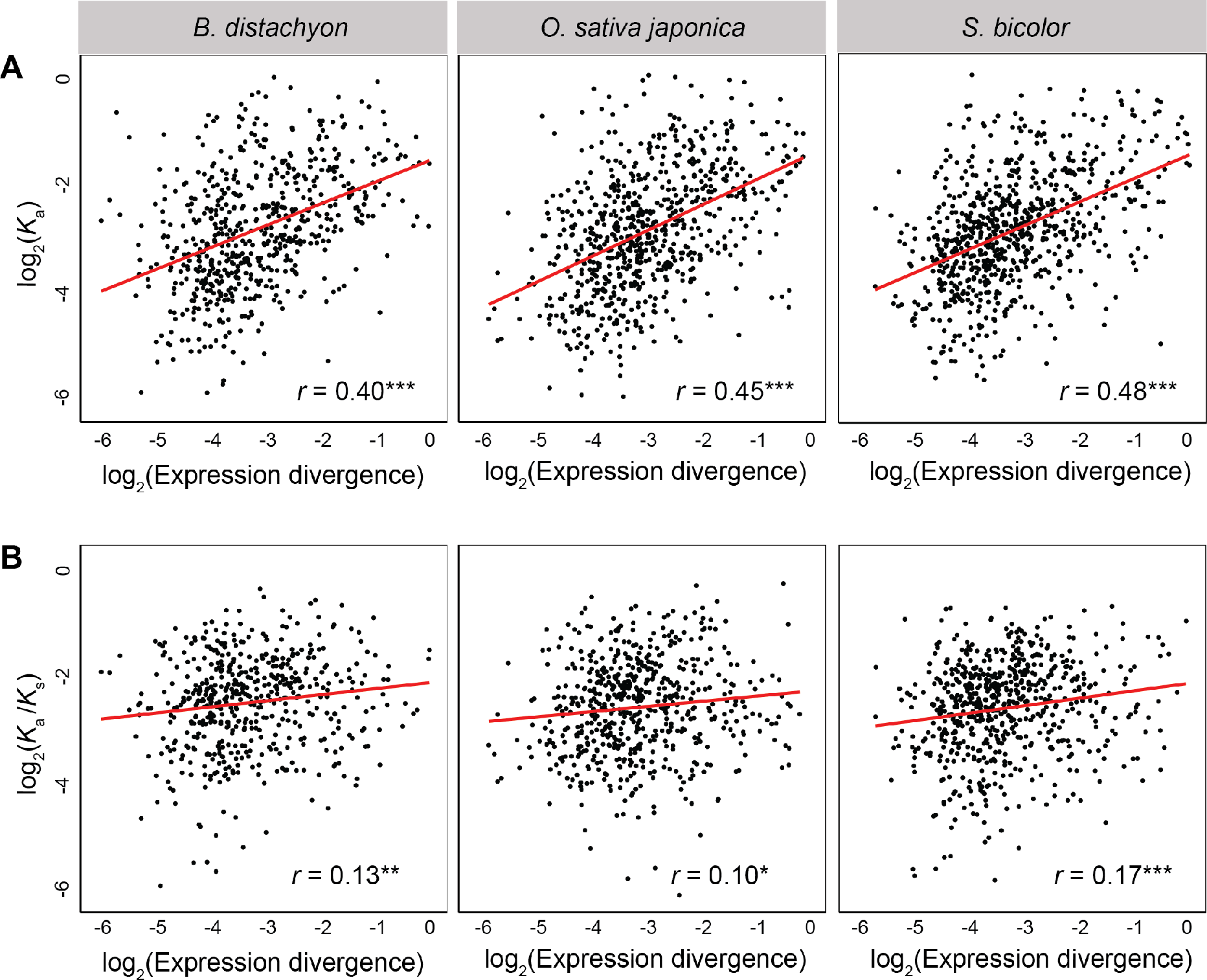
Relationship between expression and protein-coding sequence divergence rates of SSD-derived grass duplicate genes. Scatterplots showing correlations between expression divergence (Euclidian distance) and (*A*) nonsynonymous sequence divergence (*K*_a_) and (*B*) nonsynonymous/synonymous sequence divergence (*K*_a_/*K*_*S*_) rates of SSD-derived duplicate genes in *B. distachyon* (left), *O. sativa japonica* (middle), and *S. bicolor* (right). The best-fit linear regression line is shown in red, and Pearson’ s correlation coefficient (*r*) is provided at the bottom right, for each panel. \**P* < 0.05, \*\**P* < 0.01, \*\*\**P* < 0.001.

Moreover, expression divergence of SSD-derived duplicate genes is associated with increased tissue specificity in both *Drosophila* (Assis and Bachtrog 2013) and mammals (Assis and Bachtrog 2015). To assess this relationship in SSD-derived grass duplicates, we computed Pearson’ s correlation coefficients (*r*) between expression divergence (Euclidian distance) of each duplicate gene from its ancestral copy and its tissue specificity index *τ* (Yanai, et al. 2004; see Materials and Methods for details) in *B. distachyon*, *O. sativa japonica*, and *S. bicolor* (Figure 3A). Consistent with results in *Drosophila* (Assis and Bachtrog 2013) and mammals (Assis and Bachtrog 2015), there is a strong positive correlation between tissue specificity and expression divergence of SSD-derived duplicate genes in all three grass species (*r* = 0.80 − 0.87; *P* < 0.001 for all comparisons, *t* tests; see Materials and Methods for details). Thus, increased expression divergence of SSD-derived grass duplicates is associated with greater tissue specificity.

**Figure 3.**
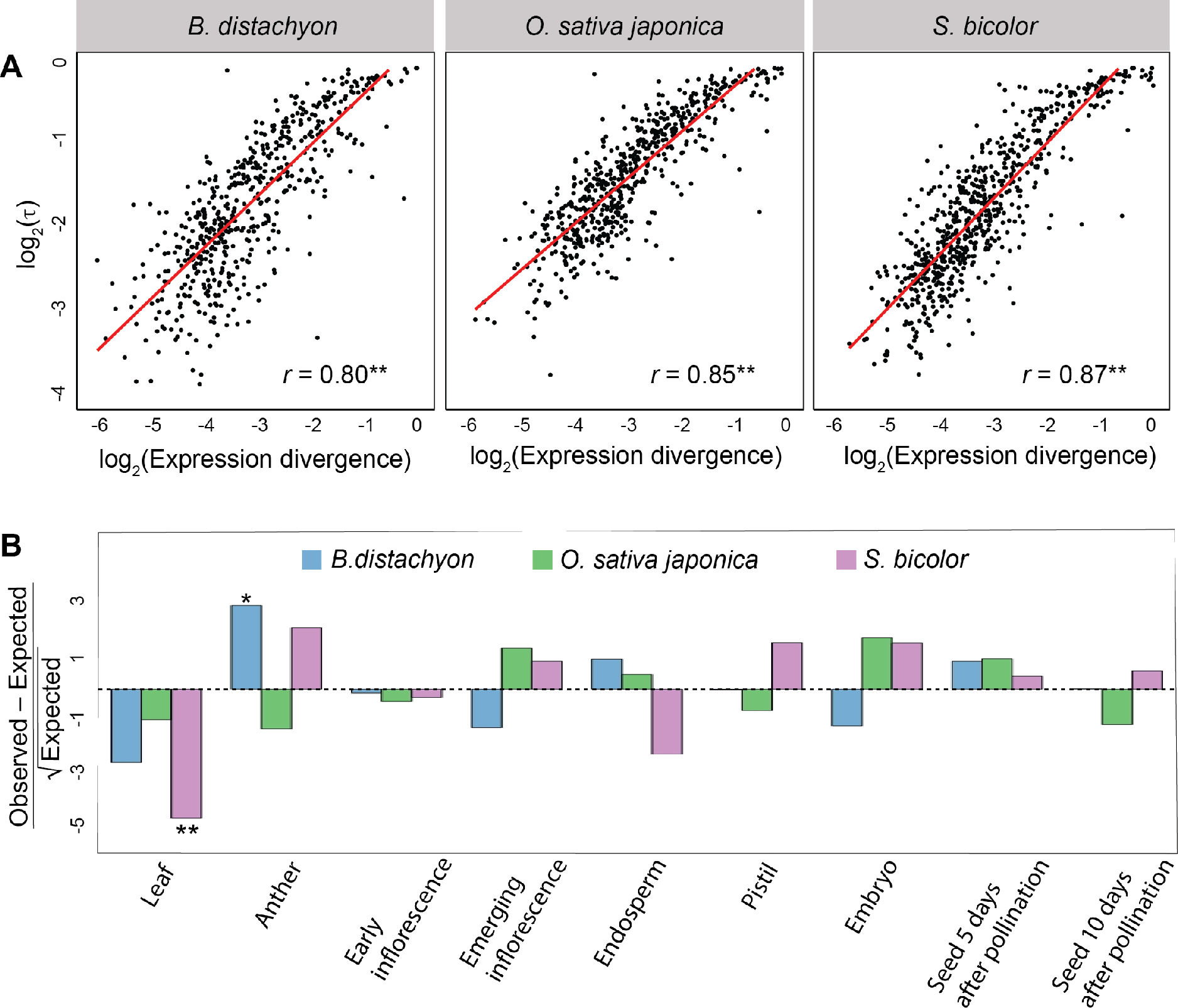
Relationship between expression divergence and tissue specificity of SSD-derived grass duplicate genes. (*A*) Scatterplot showing correlation between expression divergence (Euclidian distance) and tissue specificity (τ) of SSD-derived duplicate genes in *B. distachyon* (left), *O. sativa japonica* (middle), and *S. bicolor* (right). The best-fit linear regression line is shown in red, and Pearson’s correlation coefficient (*r*) is provided at the bottom right, for each panel. (*B*) Comparisons of observed counts of primary tissues of duplicate genes relative to those expected based on proportions of primary tissues of single-copy genes. Positive values represent overrepresentations, and negative values underrepresentations, relative to expectations. \**P* < 0.05, \*\**P* < 0.001.

Whereas SSD-derived duplicate genes in *Drosophila* are primarily testis-specific (Betrán, et al. 2002; Levine, et al. 2006b; Zhou, et al. 2008; Assis and Bachtrog 2013), those in mammals are expressed specifically in a diversity of tissues (Assis and Bachtrog 2015). Therefore, our next question was whether there are particular tissues in which SSD-derived duplicates tend to be expressed in grasses. To answer this question, we designated the tissue in which each gene has its highest expression as its primary tissue, and compared the observed primary tissues to those expected based on primary tissues of single-copy genes (Figure 3B; see Materials and Methods for details). After correcting for multiple comparisons (see Materials and Methods for details), our analysis yielded two significant findings. First, there is an underrepresentation of leaf-expressed duplicates in *S. bicolor* (*P* = 1.84×10^−6^, binomial test; see Materials and Methods for details). Because leaf is the only tissue assayed that is not related to reproduction, this result suggests that duplicates in *S. bicolor* are typically expressed in reproductive tissues. Second, we discovered an overrepresentation of anther-expressed duplicates in *B. distachyon* (*P* = 0.02, binomial test; see Materials and Methods for details). Because the anther produces pollen grains (Goldberg, et al. 1993), this result suggests that SSD-derived *B. distachyon* duplicates are involved in male-specific reproduction, as is common in many animal species (Betrán, et al. 2002; Paulding, et al. 2003; Marques, et al. 2005; Levine, et al. 2006; Vinckenbosch, et al. 2006; Zhou, et al. 2008; Assis and Bachtrog 2013, 2015). Therefore, SSD may be associated with reproduction in plants, as it is in animals.

## DISCUSSION

Despite the abundance of duplicate genes in angiosperms, and their prominent roles in evolution (Hilu 1993; Dubcovsky and Dvorak 2007; Flagel and Wendel 2009; Van de Peer, et al. 2009; Meyer, et al. 2012; Salman-Minkov, et al. 2016), their paths from genetic redundancy to functional divergence and longterm retention remain unclear. Studies in several animal species have uncovered evidence of rapid and asymmetric sequence and expression divergence after duplication that is consistent with natural selection (Conant and Wagner 2003; Blanc and Wolfe 2004; Kellis, et al. 2004; Li, et al. 2005; Assis and Bachtrog 2013, 2015; Jiang and Assis 2017). However, many angiosperms are unique in that they are self-pollinating, which may reduce their adaptive potentials (Nordborg 1997; Glémin 2012; Roze 2015; Hartfield, et al. 2017), and therefore hinder the evolutionary divergence of duplicate genes. Yet, largely due to the absence of approaches for assessing functional divergence after duplication until recently (Assis and Bachtrog 2013), no genome-wide studies have been performed to address how duplicate genes in angiosperms evolve and are retained over long evolutionary timescales. Further, previous studies in angiosperms have primarily focused on WGD-derived duplicates, whereas little emphasis has been placed on describing evolution after SSD. Therefore, our study represents the first genome-scale analysis of functional evolution after SSD in angiosperms.

Examination of expression profiles across nine tissues of *B. distachyon, O. sativa japonica*, and *S. bicolor* revealed that functional conservation is the primary long-term outcome of SSD in grasses. Conservation of duplicate genes can either be a product of selection for increased gene dosage or a consequence of slowed divergence due to a decreased efficiency of selection, which can be further exacerbated by nonallelic gene conversion. Either one or both of these mechanisms may hamper evolutionary divergence of duplicate genes in grasses. In particular, though our study focused on SSD, analyses of WGD often point to increased gene dosage as a mechanism for duplicate gene retention in plants (Bekaert, et al. 2011). On the other hand, levels of conservation in grasses are higher than those observed in *Drosophila* (Assis and Bachtrog 2013), consistent with their smaller *N*_e_ relative to *Drosophila* (Lynch and Conery 2003). Moreover, mammals and grasses have similar *N*_e_ (Lynch and Conery 2003), and the efficiency of selection in grasses is close to that in mammals. Therefore, the comparison among levels of conservation in *Drosophila*, mammals, and grasses provides additional support for a role of natural selection in evolution after gene duplication across diverse taxonomic groups.

Though our analysis suggests that most grass duplicates are functionally conserved, they also indicate that a large proportion of SSD-derived duplicates may have experienced functional divergence. Previous studies in *Arabidopsis thaliana* demonstrated that SSD-derived duplicates have greater sequence and expression divergence rates than WGD duplicates of the same age (Casneuf et al. 2006; Carretero-Paulet and Fares 2012), which may be attributed to relaxed constraint (Carretero-Paulet and Fares 2012). Therefore, it is not surprising that SSD-derived duplicates in the species considered here may have diverged functionally from their ancestral state, and it is possible that an analogous study of WGD-derived duplicates would reveal a similar trend to that observed in *A. thaliana.* Moreover, we found that expression divergence of SSD-derived grass duplicates primarily occurs asymmetrically via neofunctionalization, as has been uncovered in both *Drosophila* (Assis and Bachtrog 2013) and mammals (Assis and Bachtrog 2015). This finding is also consistent with the increased prevalence of neofunctionalization among *A. thaliana* duplicates generated by SSD (Rensing 2014). Therefore, asymmetric evolutionary divergence appears to be a common outcome of SSD in both plant and animal species. However, neofunctionalization often occurs in child copies and is associated with RNA-mediated duplication in grasses, as in *Drosophila* (Assis and Bachtrog 2013), but not in mammals (Assis and Bachtrog 2015). Further, evolutionary fates of grass duplicates are reached quickly after duplication, also consistent with findings in *Drosophila* (Assis and Bachtrog 2013), but not in mammals (Assis and Bachtrog 2015). Together, these results support the hypothesis that neofunctionalization may often occur as a byproduct of SSD itself, perhaps due to the placement of RNA-mediated duplicates in novel genomic contexts without their ancestral regulatory elements (Kaessmann, et al. 2009). Thus, aside from their slower divergence rates, the evolutionary trajectories of grass duplicates more closely mirror those of *Drosophila* (Assis and Bachtrog 2013) than mammals (Assis and Bachtrog 2015). This is somewhat surprising because the *N*_e_ of grass species are smaller than those of *Drosophila* species (Lynch and Conery 2003; Ai et al. 2012; Adugna and Bekele 2015; Stritt et al. 2017). However, in mammals, functional divergence often occurs over longer evolutionary time (Assis and Bachtrog 2015), suggesting that neofunctionalization is only biased toward child copies when it happens rapidly. This is not unexpected, given that conserved duplicates are initially redundant and, thus, the probabilities of divergence of parent and child copies over time should be equal. Therefore, this comparison further highlights the role of asymmetric duplication events, such as those that are RNA-mediated, in asymmetric divergence and child-biased neofunctionalization.

Assessment of expression divergence of SSD-derived grass duplicate genes revealed that it is positively correlated with protein-coding sequence divergence and tissue specificity. Moreover, in *B. distachyon*, we found an enrichment of duplicates highly expressed in anther, which is the tissue that produces pollen in flowering plants. This finding is consistent with those in *A. thaliana* RNA-mediated duplicates (Abdelsamad and Pecinka, 2014; Casola and Betrán, 2017) and supports the “out of the pollen” hypothesis, in which new plant genes originate from the vegetative nucleus of the mature pollen due to increased activities of transposable elements (Wu, et al. 2014). Because anther is analogous to testis in animals, our result is also synonymous with the “out of the testis” hypothesis, which posits that new genes often emerge with testis-related functions and acquire novel functions over time (Kaessmann 2010) and is supported by data in many species (Betrán, et al. 2002; Paulding, et al. 2003; Marques, et al. 2005; Levine, et al. 2006; Vinckenbosch, et al. 2006; Zhou, et al. 2008; Assis and Bachtrog 2013, 2015). Several hypotheses have been proposed to explain the male-biased origin of new genes, including increased mutation rates due to greater numbers of germline cell divisions in male tissues (Shimmin, et al. 1993), positive selection due to sexual selection (Pröschel, et al. 2006; Ellegren and Parsch 2007), and relaxed negative selection due to reduced functional pleiotropy (Ellegren and Parsch 2007; Gershoni and Pietrokovski 2014; Harrison, et al. 2015). However, as in animals (*e.g.*, Kaessmann 2010; Assis and Bachtrog 2013, 2015), any of these proposed mechanisms may contribute to the male-biased origin of duplicate genes in grasses. In particular, the increased mutation rate hypothesis (Shimmin, et al. 1993) is consistent with more cell divisions during pollen than ovule production in grasses (Filatov and Charlesworth 2002; Whittle and Johnston 2002), positive selection (Pröschel, et al. 2006; Ellegren and Parsch 2007) with the positive correlation between expression divergence and protein-coding sequence divergence of duplicates (Figure 2), and negative selection (Ellegren and Parsch 2007; Gershoni and Pietrokovski 2014; Harrison, et al. 2015) with the positive correlation between expression divergence and tissue specificity of duplicates (Figure 3A). Therefore, comparison of our findings in grasses to those in diverse animal species (Betrán, et al. 2002; Paulding, et al. 2003; Marques, et al. 2005; Levine, et al. 2006; Vinckenbosch, et al. 2006; Zhou, et al. 2008; Assis and Bachtrog 2013; Assis and Bachtrog 2014; Assis and Bachtrog 2015; Jiang and Assis 2017) highlights a universal role for gene duplication in the origin of male-specific phenotypes across plant and animal kingdoms.

## MATERIALS AND METHODS

### Identification of single-copy and duplicate genes

Reference genome annotation and sequence data from *B. distachyon* (version 1.2) (Vogel, et al. 2010), *O. sativa japonica* (version 1.0) (Sasaki, et al. 2005), and *S. bicolor* (version 1.4) (Paterson, et al. 2009), as well as a table of gene family sizes for 16 monocots, were downloaded from PLAZA 3.0 (Proost, et al. 2014) at https://bioinformatics.psb.ugent.be/plaza/. Gene families consisting of one copy in *B. distachyon, O. sativa japonica*, and *S. bicolor* were considered as single-copy genes. In total, there are 5,132 single-copy genes annotated in *B. distachyon*, 11,672 single-copy genes annotated in *O. sativa japonica*, and 6,724 single-copy genes annotated in *S. bicolor*. Removal of lowly-expressed genes (see *Sequence and expression analyses)* yielded 4,769 single-copy genes in *B. distachyon*, 5,439 single-copy genes in *O. sativa japonica* and 5,976 single-copy genes in *S. bicolor* that we used in tissue enrichment test. There are 3,466 annotated 1:1 orthologs in *B. distachyon* and *O. sativa japonica*, 3,166 annotated 1:1 orthologs in *B. distachyon* and *S. bicolor* and 3,154 annotated 1:1 orthologs in *O. sativa japonica* and *S. bicolor.* Removal of lowly-expressed genes (see *Sequence and expression analyses*) yielded 3,269 1:1 orthologs in *B. distachyon* and *O. sativa japonica*, 3,024 1:1 orthologs in *B. distachyon* and *S. bicolor* and 3,015 1:1 orthologs in *O. sativa japonica* and *S. bicolor*.

To identify pairs of duplicate genes that arose via SSD along designated branches shown in Figure 1 (full tree depicted in Figure S1), we used the maximum-likelihood method Count (Csűös 2010) to estimate rates of duplications and losses along the monocot phylogeny downloaded from PLAZA 3.0 (Proost, et al. 2014) and perform asymmetric Wagner parsimony using these rates (Swofford and Maddison 1987). In total, this approach yielded 391 pairs of duplicate genes that arose along the *B. distachyon* lineage, 478 pairs of duplicate genes that arose along the *O. sativa japonica* lineage, and 462 pairs of duplicate genes that arose along the *S. bicolor* lineage. After removing lowly-expressed genes (see *Sequence and expression analyses*), we obtained 272 pairs of *B. distachyon* duplicates, 289 pairs of *O. sativa japonica* duplicates, and 340 pairs of *S. bicolor* duplicates (see Figure 1). To assess directionality of duplications and assign parent and child copies, we used tables of orthologs from OrthoMCL (Li, et al. 2003), TribeMCL (Enright, et al. 2002), and i-ADHrRE (Fostier, et al. 2011) that were downloaded from the PLAZA 3.0 database (Proost, et al. 2014). When orthology predictions from all three methods were available yet conflicting, we applied a majority-voting scheme to infer the most likely orthologs. When predictions from only two methods were available and conflicting, we prioritized OrthoMCL orthologs above all others, and i-ADHrRE above TribeMCL.

### Sequence and expression analyses

We performed all sequence alignments between duplicates and ancestral single-copy genes using MACSE 1.0 (Ranwez, et al. 2011), which accounts for frameshifts and stop codons. We estimated *K*_a_ and *K*_a_/*K*_s_ ratios using the codeml package in PAML 4.0 (Yang 2007) with runmode = −2, model = 0, and NSsites = 0. To avoid saturation at synonymous sites, we only considered *K*_s_ < 3. Tables containing expression abundances estimated in transcripts per million (TPM) from RNA-seq data of protein-coding genes in nine tissues (leaf, anther, endosperm, early inflorescence, emerging inflorescence, pistil, embryo, seed 5 days after pollination, and seed 10 days after pollination) (Davidson, et al. 2012) of *Brachypodium distachyon, Oryza sativa japonica*, and *Sorghum bicolor* were downloaded from Expression Atlas at https://www.ebi.ac.uk/gxa/home. These RNA-seq data were quantified with HTSeq 0.6 (Anders, et al. 2015), which only counts reads that unambiguously map to a single gene, thereby minimizing the probability of incorrect mapping between duplicate gene copies. Data were then log-transformed, and genes with log_2_(TPM + 1) < 1 in all nine tissues were removed. We estimated the expression breadth of each gene with the tissue specificity index τ (Yanai, et al. 2004), which is defined as 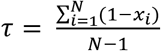, where *x*_*i*_ represents the expression level in the *i*^*th*^ tissue normalized by the maximal expression value. The range of τ is from 0 to 1, with larger τ signifying greater tissue specificity.

We classified retention mechanisms of duplicate genes in our dataset using the CDROM R package (Perry and Assis 2016), which implements Assis and Bachtrog’ s phylogenetic approach (Assis and Bachtrog 2013). In particular, CDROM takes as input tables of expression measurements for multiple conditions in two sister species, lists of orthologous single-copy genes in the two sisters, and a list of parent and child duplicate gene pairs in one sister and their ancestral genes in the second sister. We used *B. distachyon* as the sister species to *O. sativa* and *S. bicolor* and applied CDROM to the RNA-seq data described above, which consists of log-transformed TPMs for genes in nine tissues of *B. distachyon, O. sativa japonica*, and *S. bicolor* (Davidson, et al. 2012). CDROM first calculates Euclidian distances between expression profiles of orthologous single-copy genes (*E*_*S*1,*S*2_), expression profiles of parent and child duplicate genes and the ancestral gene (*E*_*P, A*_ and *E*_*C,A*_), and combined expression profiles of both duplicate genes and the ancestral gene (*E*_*P*+*CA*_). Next, it uses a user-specific cutoff for *E*_*S*1,*S*2_ (*E*_*div*_) to classify retention mechanisms of duplicates. Specifically, duplicates with *E*_*P,A*_ ≤ *E*_*div*_ and *E*_*C,A*_ ≤ *E*_*div*_ are classified as functionally conserved; those with either *E*_*P,A*_ ≤ *E*_*div*_ and *E*_*C,A*_ > *E*_*div*_ or *E*_*C,A*_ ≤ *E*_*div*_ and *E*_*P,A*_ > *E*_*div*_ as neofunctionalized; those with *E*_*P,A*_ > *E*_*div*_, *E*_*C,A*_ > *E*_*div*_, and *E*_*P+CA*_ ≤ *E*_*div*_ as subfunctionalized, and those with *E*_*PA*_ > *E*_*div*_, *E*_*C,A*_ > *E*_*div*_ and *E*_*P+C,A*_ > *E*_*div*_ as specialized. We used distributions of Euclidian distances between gene expression profiles to choose *E*_*div*_ for each species (Figure S2).

### Determination of DNA-and RNA-mediated duplication mechanisms

Exon counts for parent and child duplicates were obtained from genome annotation files (*B. distachyon* version 1.2 (Vogel, et al. 2010), *O. sativa japonica* version 1.0 (Sasaki et al. 2005), and *S. bicolor* version 1.4 (Paterson, et al. 2009)) downloaded from the PLAZA 3.0 database (Proost, et al. 2014). The child was considered as arising through DNA-mediated duplication when the parent and child copies both have multiple exons, and through RNA-mediated duplication when the parent copy has multiple exons and the child copy has one exon. When both the parent and child have one exon, the mechanism was considered to be unknown (43 pairs in *Brachypodium distachyon*, 39 in *Oryza sativa japonica* and 50 in *Sorghum bicolor*). Genes with unknown duplication mechanisms genes were not used in the analysis presented in Table 2.

### Statistical analyses

We performed all statistical analyses in the R software environment (R Core Team 2013). *χ*^2^ tests were used to compare observed and expected DNA-and RNA-mediated duplicates retained through different mechanisms (Table 2), as well as observed and expected retention mechanisms of duplicates in different age groups (Tables S1-3). Expected counts of DNA-and RNA-mediate duplicates were obtained by multiplying the number of duplicates retained by each mechanism by total proportions of DNA-and RNA-mediated duplicates, respectively. Expected counts of retention mechanisms of duplicates in different age groups were obtained by multiplying the number of duplicates retained by each mechanism by total proportions of duplicates in different age groups. Significance of Pearson s correlation coefficients depicted in Figures 2 were assessed via Student’ s *t* tests. Two-tailed binomial tests were implemented to compare observed counts of highest-expressed duplicates relative to their expected probabilities. Each binomial test was performed by setting the number of trials as the total number of duplicates, the number of successes as the number of highest-expressed duplicates in the tissue of interest, and the probability of success as the frequency of single-copy genes in the tissue of interest. P-values from binomial tests were Bonferroni-adjusted to correct for the nine comparisons performed.

## Supporting information

## REFERENCES

Abdelsamad A, Pecinka A. 2014. Pollen-specific activation of *Arabidopsis* retrogenes is associated with global transcriptional reprogramming. The Plant Cell 1:114.

Adugna A, Bekele E. 2015. Assessment of recent bottlenecks and estimation of effective population size in the Ethiopian wild sorghum using simple sequence repeat allele diversity and mutation models. Plant Genet Resour 13:274–281.

Ai B, Wang ZS, Ge S. 2012. Genome size is not correlated with effective population size in the *Oryza* species. Evolution 66:3302–3310.

Aklilu BB, Soderquist RS, Culligan KM. 2013. Genetic analysis of the Replication Protein A large subunit family in *Arabidopsis* reveals unique and overlapping roles in DNA repair, meiosis and DNA replication. Nucleic Acids Res 42:3104–3118.

Anders S, Pyl PT, Huber W. 2015. HTSeq—a Python framework to work with high-throughput sequencing data. Bioinformatics 31:166–169.

Assis R. 2014. *Drosophila* duplicate genes evolve new functions on the fly. Fly 8:91–94.

Assis R, Bachtrog D. 2013. Neofunctionalization of young duplicate genes in *Drosophila*. Proc Natl Acad Sci USA 110:17409–17414.

Assis R, Bachtrog D. 2015. Rapid divergence and diversification of mammalian duplicate gene functions. BMC Evol Biol 15:138.

Beachell H, Adair CR, Jodon N, Davis L, Jones JW. 1938. Extent of natural crossing in rice. J Am Soc Agronomy.

Bekaert M, Edger PP, Pires JC, Conant GC. 2011. Two-phase resolution of polyploidy in the *Arabidopsis* metabolic network gives rise to relative and absolute dosage constraints. The Plant Cell 23:1719–1728.

Bennetzen JL, Ma J, Devos KM. 2005. Mechanisms of recent genome size variation in flowering plants. Annals Bot 95:127–132.

Bennetzen JL. 2007. Patterns of grass genome evolution. Curr Opin Plant Biol 10:176–181.

Betrán E, Thornton K, Long M. 2002. Retroposed new genes out of the X in *Drosophila*. GenomeRes 12:1854–1859.

Blanc G, Wolfe KH. 2004. Functional divergence of duplicated genes formed by polyploidy during *Arabidopsis* evolution. The Plant Cell 16:1679–1691.

Bowers JE, Arias MA, Asher R, Avise JA, Ball RT, Brewer GA, Buss RW, Chen AH, Edwards TM, Estill JC et al. 2005. Comparative physical mapping links conservation of microsynteny to chromosome structure and recombination in grasses. Proc Natl Acad Sci USA, 102:13206–13211.

Carretero-Paulet L, Fares MA. 2012. Evolutionary dynamics and functional specialization of plant paralogs formed by whole and small-scale genome duplications. Mol Biol and Evol 9:3541–3551.

Casneuf T, De Bodt S, Raes J, Maere S, Van de Peer Y. 2006. Nonrandom divergence of gene expression following gene and genome duplications in the flowering plant *Arabidopsis* thaliana. Genome Biol 7:R13.

Casola C, Betrán E. 2017. The genomic impact of gene retrocopies: what have we learned from comparative genomics, population genomics, and transcriptomic analyses?. Genome Biol Evol 9:1351–1373.

Charlesworth B. 2009. Effective population size and patterns of molecular evolution and variation. Nat Rev Genet 10:195–205.

Chau LM, Goodisman MA. 2017. Gene duplication and the evolution of phenotypic diversity in insect societies. Evolution 71:2871–2884.

Conant GC, Wagner A. 2003. Asymmetric sequence divergence of duplicate genes. Genome Res 13:2052–2058.

Conant GC, Wagner A. 2004. Duplicate genes and robustness to transient gene knock-downs in *Caenorhabditis elegans*. Proc R Soc Lond [Biol] 271:89–96.

Csűös M. 2010. Count: evolutionary analysis of phylogenetic profiles with parsimony and likelihood. Bioinformatics 26:1910–1912.

Cui L, Wall PK, Leebens-Mack JH, Lindsay BG, Soltis DE, Doyle JJ, Soltis PS, Carlson JE, Arumuganathan K, Barakat A. 2006. Widespread genome duplications throughout the history of flowering plants. Genome Res 16:738–749.

Davidson RM, Gowda M, Moghe G, Lin H, Vaillancourt B, Shiu SH, Jiang N, Robin Buell C. 2012. Comparative transcriptomics of three Poaceae species reveals patterns of gene expression evolution. The Plant Journal 71:492–502.

Dje Y, Heuertz M, Lefebvre C, Vekemans X. 2000. Assessment of genetic diversity within and among germplasm accessions in cultivated sorghum using microsatellite markers. Theor Appl Genet 100:918–925.

Duarte JM, Cui L, Wall PK, Zhang Q, Zhang X, Leebens-Mack J, Ma H, Altman N, DePamphilis CW. 2005. Expression pattern shifts following duplication indicative of subfunctionalization and neofunctionalization in regulatory genes of *Arabidopsis*. Mol Biol Evol 23:469–478.

Dubcovsky J, Dvorak J. 2007. Genome plasticity a key factor in the success of polyploid wheat under domestication. Science 316:1862–1866.

Ellegren H, Parsch J. 2007. The evolution of sex-biased genes and sex-biased gene expression. Nat Rev Genet. 8:689–698.

Enright AJ, Van Dongen S, Ouzounis CA. 2002. An efficient algorithm for large-scale detection of protein families. Nucleic Acid Res 30:1575–1584.

Flagel LE, Wendel JF. 2009. Gene duplication and evolutionary novelty in plants. New Phytologist 183:557–564.

Flavell R, Bennett M, Smith J, Smith D. 1974. Genome size and the proportion of repeated nucleotide sequence DNA in plants. Biochem Genet 12:257–269.

Force A, Lynch M, Pickett FB, Amores A, Yan Y-l, Postlethwait J. 1999. Preservation of duplicate genes by complementary, degenerative mutations. Genetics 151:1531–1545.

Fostier J, Proost S, Dhoedt B, Saeys Y, Demeester P, Van de Peer Y, Vandepoele K. 2011. A greedy, graph-based algorithm for the alignment of multiple homologous gene lists. Bioinformatics 27:749–756.

Freeling M. 2009. Bias in plant gene content following different sorts of duplication: tandem, whole-genome, segmental, or by transposition. Annual Rev Plant Biol 60:433–453.

Gershoni M, Pietrokovski S. 2014. Reduced selection and accumulation of deleterious mutations in genes exclusively expressed in men. Nat Commun 5:4438.

Glémin S. 2012. Extinction and fixation times with dominance and inbreeding. Theore Popul Biol 81:310316.

Goldberg RB, Beals TP, Sanders PM. 1993. Anther development: basic principles and practical applications. The Plant Cell 5:1217.

Gordon SP, Priest H, Des Marais DL, Schackwitz W, Figueroa M, Martin J, Bragg JN, Tyler L, Lee CR, Bryant D. 2014. Genome diversity in *Brachypodium distachyon*: deep sequencing of highly diverse inbred lines. The Plant Journal 79:361–374.

Gossmann TI, Keightley PD, Eyre-Walker A. 2012. The effect of variation in the effective population size on the rate of adaptive molecular evolution in eukaryotes. Genome Biol Evol 4:658–667.

Gu Z, Nicolae D, Lu HH, Li W-H. 2002. Rapid divergence in expression between duplicate genes inferred from microarray data. Trends Genet 18:609–613.

Gu Z, Rifkin SA, White KP, Li WH. 2004. Duplicate genes increase gene expression diversity within and between species. Nature Genet 36:577.

Hakes L, Pinney JW, Lovell SC, Oliver SG, Robertson DL. 2007. All duplicates are not equal: the difference between small-scale and genome duplication. Genome Biol 8:R209.

Han MV, Hahn MW. 2009. Identifying parent-daughter relationships among duplicated genes. Biocomputing:114–125.

Harrison PW, Wright AE, Zimmer F, Dean R, Montgomery SH, Pointer MA, Mank JE. 2015. Sexual selection drives evolution and rapid turnover of male gene expression. Proc Natl Acad Sci USA. 112:4393–4398.

Hartfield M, Bataillon T, Glémin S. 2017. The evolutionary interplay between adaptation and selfFertilization. Trends Genet 33:420–431.

He X, Zhang J. 2005. Rapid subfunctionalization accompanied by prolonged and substantial neofunctionalization in duplicate gene evolution. Genetics 169:1157–1164.

Hilu K. 1993. Polyploidy and the evolution of domesticated plants. Am J Bot 1494–1499.

International Brachypodium Initiative. 2010. Genome sequencing and analysis of the model grass *Brachypodium* distachyon. Nature 463:763–768.

Jiang X, Assis R. 2017. Natural selection drives rapid functional evolution of young *Drosophila* duplicate genes. Mol Biol Evol 34:3089–3098.

Jiao Y, Wickett NJ, Ayyampalayam S, Chanderbali AS, Landherr L, Ralph PE, Tomsho LP, Hu Y, Liang H, Soltis PS, Soltis DE, et al. 2011. Ancestral polyploidy in seed plants and angiosperms. Nature. 473(7345):97.

Kaessmann H. 2010. Origins, evolution, and phenotypic impact of new genes. Genome Res 20:13131326.

Kaessmann H, Vinckenbosch N, Long M. 2009. RNA-based gene duplication: mechanistic and evolutionary insights. Nat Rev Genet 10:19.

Kellis M, Birren BW, Lander ES. 2004. Proof and evolutionary analysis of ancient genome duplication in the yeast *Saccharomyces cerevisiae*. Nature 428:617–624.

Kimura M. 1983. The neutral theory of molecular evolution: Cambridge University Press.

Levine MT, Jones CD, Kern AD, Lindfors HA, Begun DJ. 2006a. Novel genes derived from noncoding DNA in *Drosophila melanogaster* are frequently X-linked and exhibit testis-biased expression. Proc Natl Acad Sci USA 103:9935–9939.

Levine MT, Jones CD, Kern AD, Lindfors HA, Begun DJ. 2006b. Novel genes derived from noncoding DNA in *Drosophila melanogaster* are frequently X-linked and exhibit testis-biased expression. Proc Natl Acad Sci USA 103:9935–9939.

Li L, Stoeckert CJ, Roos DS. 2003. OrthoMCL: identification of ortholog groups for eukaryotic genomes. Genome Res 13:2178–2189.

Li W-H, Yang J, Gu X. 2005. Expression divergence between duplicate genes. Trends Genet 21:602–607.

Lockton S, Gaut BS. 2005. Plant conserved non-coding sequences and paralogue evolution. Trends Genet 21:60–65.

Lynch M, Conery JS. 2003. The origins of genome complexity. Science 302:1401–1404.

Ma Y, Wang J, Zhong Y, Geng F, Cramer GR, Cheng Z-MM. 2015. Subfunctionalization of cation/proton antiporter 1 genes in grapevine in response to salt stress in different organs. Hort Res 2:15031.

Makova KD, Li W-H. 2003. Divergence in the spatial pattern of gene expression between human duplicate genes. Genome Res 13:1638–1645.

Marcussen T, Oxelman B, Skog A, Jakobsen KS. 2010. Evolution of plant RNA polymerase IV/V genes: evidence of subneofunctionalization of duplicated *NRPD2/NRPE2*-like paralogs in *Viola* (Violaceae). BMC Evol Biol 10:45.

Maere S, De Bodt S, Raes J, Casneuf T, Van Montagu M, Kuiper M, Van de Peer Y. 2005. Modeling gene and genome duplications in eukaryotes. Proc Natl Acad Sci USA 10:5454–5459.

Maere S, Van de Peer Y. 2010. Duplicate retention after small-and large-scale duplications. In K Dittmar, DA Liberles, eds, Evolution after Gene Duplication. John Wiley & Sons, Hoboken, NJ, pp 31–56

Marques AC, Dupanloup I, Vinckenbosch N, Reymond A, Kaessmann H. 2005. Emergence of young human genes after a burst of retroposition in primates. PLoS Biol 3:e357.

Meyer RS, DuVal AE, Jensen HR. 2012. Patterns and processes in crop domestication: an historical review and quantitative analysis of 203 global food crops. New Phytologist 196:29–48.

Nordborg M. 1997. Structured coalescent processes on different time scales. Genetics 146:1501–1514.

Ohno S. 1970. Evolution by gene duplication: Springer Science & Business Media.

Panchy, Nicholas, Melissa Lehti-Shiu, and Shin-Han Shiu. 2016. Evolution of gene duplication in plants. Plant Physiol 171: 2294–2316.

Paterson A, Bowers J, Bruggmann R, Dubchak I, Grimwood J, Gundlach H, Haberer G, Hellsten U, Mistros T, Poliakov A, el al. 2009. The *Sorghum bicolor* genome and the diversification of grasses. Nature 457:551–556.

Paterson A, Bowers J, Chapman B. 2004. Ancient polyploidization predating divergence of the cereals, and its consequences for comparative genomics. Proc Natl Acad Sci USA 101:9903–9908.

Paulding CA, Ruvolo M, Haber DA. 2003. The Tre2 (USP6) oncogene is a hominoid-specific gene. Proc Natl Acad Sci USA 100:2507–2511.

Perry BR, Assis R. 2016. CDROM: Classification of duplicate gene retention mechanisms. BMC Evol Biol 16:82.

Proost S, Van Bel M, Vaneechoutte D, Van de Peer Y, Inzé D, Mueller-Roeber B, Vandepoele K. 2014. PLAZA 3.0: an access point for plant comparative genomics. Nucleic Acid Res 43:D974–D981.

Pröschel M, Zhang Z, Parsch J. 2006 Widespread adaptive evolution of *Drosophila* genes with sex-biased expression. Genetics 174: 893–900

R Core Team. 2013. R: A language and environment for statistical computing. R Foundation for Statistical Computing, Vienna, Austria.

Ranwez V, Harispe S, Delsuc F, Douzery EJ. 2011. MACSE: Multiple Alignment of Coding SEquences accounting for frameshifts and stop codons. PLoS One 6:e22594.

Rastogi S, Liberles DA. 2005. Subfunctionalization of duplicated genes as a transition state to neofunctionalization. BMC Evol Biol 5:28.

Rensing SA. 2014. Gene duplication as a driver of plant morphogenetic evolution. Curr Opin Plant Biol 17:43–48.

Seoighe C, Gehring C. Genome duplication led to highly selective expansion of the *Arabidopsis* thaliana proteome. Trends in Genetics. 2004 Oct 1;20(10):461–4.

Roze D. 2015. Effects of interference between selected loci on the mutation load, inbreeding depression, and heterosis. Genetics 201:745–757.

Salman-Minkov A, Sabath N, Mayrose I. 2016. Whole-genome duplication as a key factor in crop domestication. Nature plants 2:16115.

Sasaki T. 2005. The map-based sequence of the rice genome. Nature 436:793.

Shimmin LC, Chang BH, Li WH. 1993. Male-driven evolution of DNA sequences. Nature 362:745 – 747

Stoltzfus A. 1999. On the possibility of constructive neutral evolution. J Mol Evol 49:169–181.

Stritt C, Gordon SP, Wicker T, Vogel JP, Roulin AC. 2017. Recent activity in expanding populations and purifying selection have shaped transposable element landscapes across natural accessions of the Mediterranean grass *Brachypodium distachyon*. Genome Biol Evol 10:304–318.

Swofford DL, Maddison WP. 1987. Reconstructing ancestral character states under Wagner parsimony. Math Biosci 87:199–229.

Throude M, Bolot S, Bosio M, Pont C, Sarda X, Quraishi UM, Bourgis F, Lessard P, Rogowsky P, Ghesquiere A. 2009. Structure and expression analysis of rice paleo duplications. Nucleic Acids Res 37:1248–1259.

Van de Peer Y, Maere S, Meyer A. 2009. The evolutionary significance of ancient genome duplications. Nat Rev Genet 10:725.

Vinckenbosch N, Dupanloup I, Kaessmann H. 2006. Evolutionary fate of retroposed gene copies in the human genome. Proc Natl Acad Sci USA 103:3220–3225.

Wu D-D, Wang X, Li Y, Zeng L, Irwin DM, Zhang Y-P. 2014. “Out of pollen” hypothesis for origin of new genes in flowering plants: study from *Arabidopsis thaliana*. Genome Biol Evol 6:2822–2829.

Yanai I, Benjamin H, Shmoish M, Chalifa-Caspi V, Shklar M, Ophir R, Bar-Even A, Horn-Saban S, Safran M, Domany E. 2004. Genome-wide midrange transcription profiles reveal expression level relationships in human tissue specification. Bioinformatics 21:650–659.

Yang Z. 2007. PAML 4: a program package for phylogenetic analysis by maximum likelihood. Mol Biol Evol 24:1568–1591.

Zhang J. 2003. Evolution by gene duplication: an update. Trend Ecol Evol 18:292–298.

Zhang S, Zhang J-S, Zhao J, He C. 2015. Distinct subfunctionalization and neofunctionalization of the B-class MADS-box genes in *Physalis floridana*. Planta 241:387–402.

Zhang Z, Gu J, Gu X. 2004. How much expression divergence after yeast gene duplication could be explained by regulatory motif evolution? Trends Genet 20:403–407.

Zhou Q, Zhang G, Zhang Y, Xu S, Zhao R, Zhan Z, Li X, Ding Y, Yang S, Wang W. 2008. On the origin of new genes in *Drosophila*. Genome Res 18:1446–1455.

