## Supplementary material for "Rapid functional divergence of grass duplicate genes"

**Table S1. Observed (expected) counts of *B. distachyon* retention mechanisms by SSD age**

| | Conservation | Neofunctionalization (child) | Neofunctionalization (parent) | Specialization | $P$ |
|---|---|---|---|---|---|
| *H. vulgare* | 113 (115.05) | 31 (28.93) | 11 (11.44) | 28 (27.58) | 0.98 |
| *O. sativa japonica* | 36 (37.09) | 8 (9.33) | 5 (3.69) | 10 (8.89) | 0.85 |
| *S. bicolor* | 22 (18.86) | 4 (4.74) | 1 (1.88) | 3 (4.52) | 0.69 |

Duplication ages are given as divergence times from Figure 1 and are listed from youngest to oldest.

**Table S2. Observed (expected) counts of *O. sativa japonica* retention mechanisms by SSD age**

| | Conservation | Neofunctionalization (child) | Neofunctionalization (parent) | Specialization | $P$ |
|---|---|---|---|---|---|
| *O. sativa indica* | 32 (31.82) | 13 (11.28) | 4 (4.83) | 9 (10.07) | 0.92 |
| *B. distachyon* | 108 (110.27) | 39 (39.08) | 15 (16.75) | 39(34.90) | 0.87 |
| *S. bicolor* | 18 (15.91) | 4 (5.64) | 5 (2.42) | 2 (5.03) | 0.14 |

Duplication ages are given as divergence times from Figure 1 and are listed from youngest to oldest.

**Table S3. Observed (expected) counts of *S. bicolor* retention mechanisms by SSD age**

| | Conservation | Neofunctionalization (child) | Neofunctionalization (parent) | Specialization | $P$ |
|---|---|---|---|---|---|
| *Z. mays* | 103 (101.26) | 27 (30.65) | 5 (3.72) | 22 (21.37) | 0.82 |
| *S. italica* | 75 (72.24) | 28 (21.87) | 2 (2.65) | 7 (15.24) | 0.09 |
| *B. distachyon/ O. sativa japonica* | 40 (44.50) | 11 (13.48) | 1 (1.63) | 17 (9.39) | 0.07 |

Duplication ages are given as divergence times from Figure 1 and are listed from youngest to oldest.

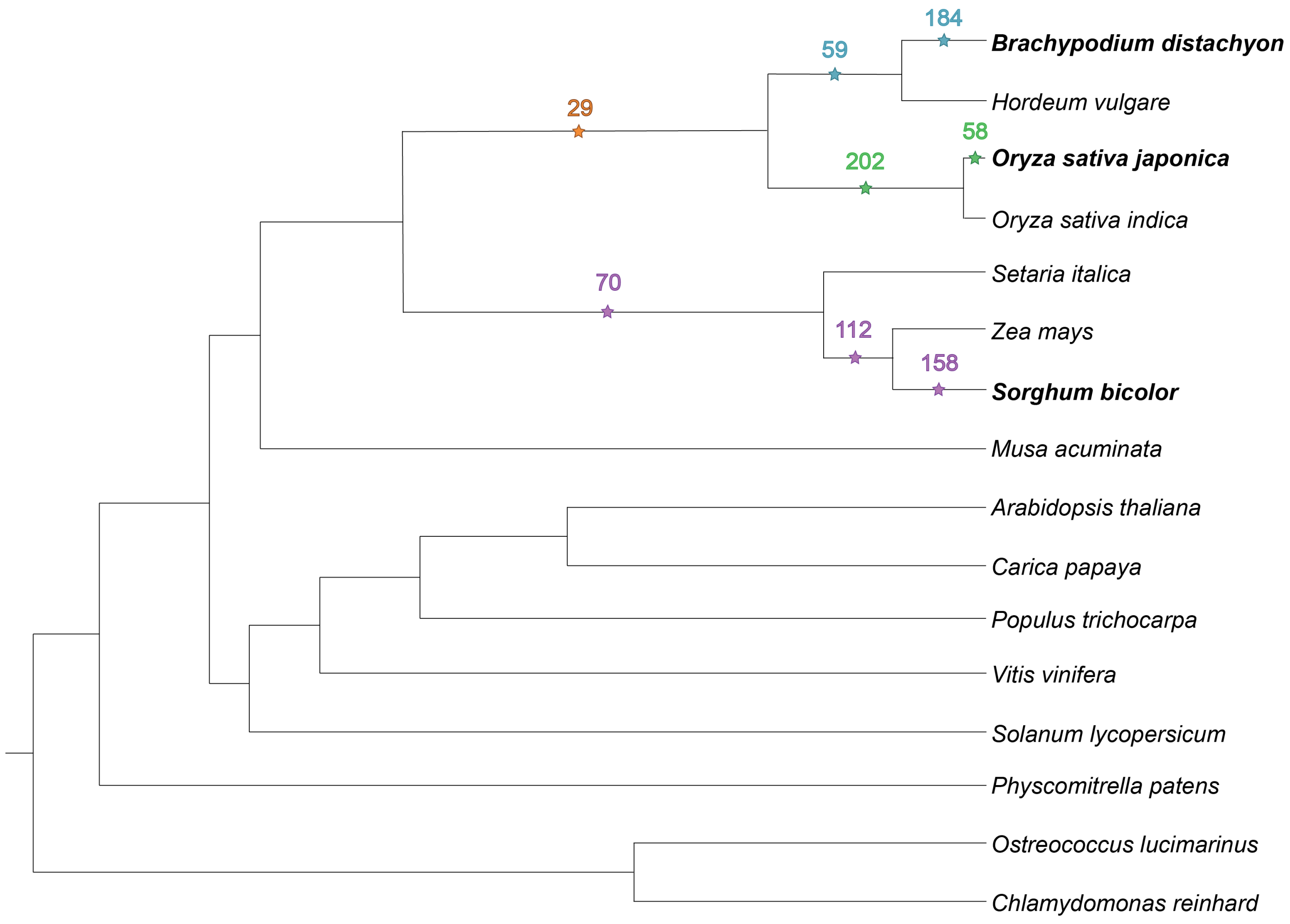


**Figure S1. Monocot phylogeny used to infer SSD events**. Numbers of duplicate gene pairs that arose via SSD along the *B. distachyon* (blue and orange stars), *O. sativa japonica* (green and orange stars), and *S. bicolor* (purple stars) lineages at specified divergence times on the monocot phylogeny.

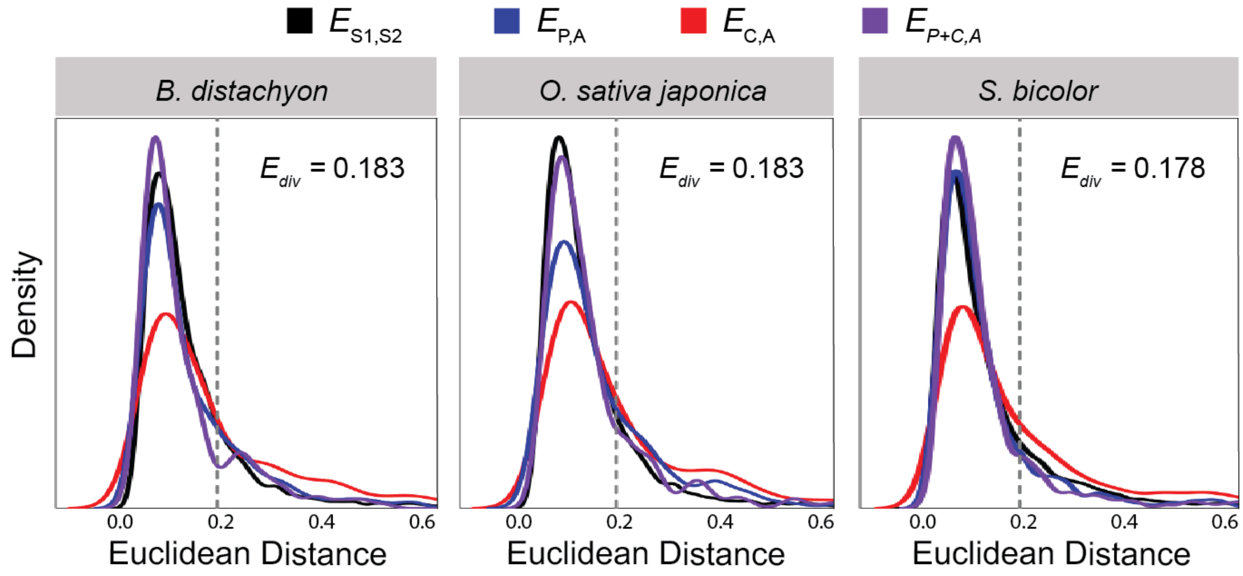


**Figure S2. Distributions of Euclidean distances between gene expression profiles in *B. distachyon* (left), *O. sativa japonica* (middle), and *S. bicolor* (right).** Distances were calculated between expression profiles of single-copy genes ($E_{\mathrm{S1,S2}}$, black), parent duplicates and ancestral genes ($E_{\mathrm{P,A}}$, blue), child duplicates and ancestral genes ($E_{\mathrm{C,A}}$, red), and parent and child duplicates combined and ancestral genes ($E_{\mathrm{P+C,A}}$, purple). Vertical dashed lines represent cutoffs ($E_{\mathrm{div}}$) used to assess expression divergence of duplicate genes in each species.
